# Metaphor - A workflow for streamlined assembly and binning of metagenomes

**DOI:** 10.1101/2023.02.09.527784

**Authors:** Vinícius W. Salazar, Babak Shaban, Maria del Mar Quiroga, Robert Turnbull, Edoardo Tescari, Vanessa Rossetto Marcelino, Heroen Verbruggen, Kim-Anh Lê Cao

## Abstract

Recent advances in bioinformatics and high-throughput sequencing have enabled the large-scale recovery of genomes from metagenomes. This has the potential to bring important insights as researchers can bypass cultivation and analyse genomes sourced directly from environmental samples. There are, however, technical challenges associated with this process, most notably the complexity of computational workflows required to process metagenomic data, which include dozens of bioinformatics software tools, each with their own set of customisable parameters that affect the final output of the workflow. At the core of these workflows are the processes of assembly - combining the short input reads into longer, contiguous fragments (contigs), and binning - clustering these contigs into individual genome bins. Both processes can be done for each sample separately or by pooling together multiple samples to leverage information from a combination of samples. Here we present Metaphor, a fully-automated workflow for genome-resolved metagenomics (GRM). Metaphor differs from existing GRM workflows by offering flexible approaches for the assembly and binning of the input data, and by combining multiple binning algorithms with a bin refinement step to achieve high quality genome bins. Moreover, Metaphor generates reports to evaluate the performance of the workflow. We showcase the functionality of Metaphor on different synthetic datasets, and the impact of available assembly and binning strategies on the final results. The workflow is freely available at https://github.com/vinisalazar/metaphor.

**Author summary:** We present Metaphor, a user-friendly, automated workflow for the recovery of genomes from metagenomes. Our tool offers flexible options for assembling and binning metagenomic contigs, that may be adjusted according to the characteristics of the input data and available computational resources, and a combination of binning algorithms, which improves the quantity and quality of resulting genome bins. We showcase the performance of Metaphor on synthetic benchmarking datasets and discuss the implication of methodological decisions regarding the strategy for assembling and binning metagenomic contigs.

## Introduction

Genome-resolved metagenomics (GRM) is a set of techniques for the recovery of genomes from high-throughput sequencing data. In recent years, applications of GRM have led to unprecedented insight into microbial diversity, ecology, and evolution, due to the recovery of (mostly uncultivated) metagenome-assembled genomes (MAGs) [1–4]. MAGs are essentially “bins” of contigs that are clustered together based on differential coverage and sequence composition; a bin is considered a MAG when it displays a high degree of completeness and a low degree of redundancy/contamination, which is usually calculated through the presence of marker genes in the bin. Advances in GRM have consistently improved the quality of recovered MAGs, and large-scale studies reconstructing and analysing thousands of MAGs have become prominent in microbiology research. Even with the inherent biases that accompany the generation of MAGs, it is evident that the benefits outweigh the risks, and researchers are increasingly in need of automated data processing methods for assembling and binning metagenomes [5]. Data pipelines that perform such experiments are inherently complex, have high computing cost, use heterogeneous data sources, have dozens of customisable parameters, and depend on several specialised bioinformatics software [6, 7].

An additional domain-specific challenge for GRM studies is the strategy used for assembling and binning each sequenced sample. Data (raw reads generated by the sequencer) originating from multiple samples may be assembled separately or pooled together, depending whether they come from the same population, specimen, or environment. This results in either a set of contigs for each sample or a ‘coassembly’ of the pooled samples. Similarly, in the metagenome binning step, where contigs are clustered into genome bins, one may do this individually for each set of assembled contigs, or by pooling together contigs from multiple samples and then mapping each individual sample to this catalogue of contigs (‘cobinning’) [8]. The latter approach allows binning algorithms to account for differential coverage of contigs across samples, enriching the information available for clustering. The chosen strategy for assembly and binning may have important consequences for the final results, *i.e*., the quality of the assembly and of the recovered bins [8]. It is hypothesised that pooled assembly and binning may lead to improved results when analysing communities with high genetic diversity, and to poorer results when there is a high level of intraspecies/strain-level diversity [9],

Here we present Metaphor, an automated and flexible workflow for the assembly and binning of metagenomes, which recovers prokaryotic genomes from metagenomes efficiently and with high sensitivity, and provides taxonomic and functional abundance data for quantitative metagenome analyses. Our software advances existing metagenomic pipelines by combining two core features: the usage of multiple binning software along with a binning refinement step, and the possibility of defining groups for assembly and binning of samples. This effectively allows scaling Metaphor to process multiple datasets in a single execution, performing assembly and binning in separate batches for each dataset, and avoiding the need for repeated executions with different input datasets. The workflow includes native functionality for downstream integration with ‘omics statistical toolkits [10, 11], so that abundance data can be easily imported into these tools, and with the Anvi’o [12] platform, which allows importing the collections of bins generated by Metaphor along with contig coverage data.. Metaphor generates detailed performance metrics at the end of each module of the workflow to provide users with a high-level summary of their analysis, and has been designed to be user-friendly, portable, and flexible, as users can choose between different strategies for assembly and binning. We demonstrate its functionality using different synthetic datasets and discuss how these different strategies can impact data analyses in terms of quality of the resulting assembly and genome bins.

## Design and Implementation

Metaphor stands out from existing GRM pipelines by offering flexible options for assembly and binning combined with multiple binning software and a binning refinement step. See Table 1 for a comparison of Metaphor’s features with other state-of-the-art GRM workflows. The workflow is implemented with Snakemake [13], a widely-used scientific workflow management system. In each module, computing steps (called “rules” by Snakemake) consist of both third-party bioinformatics software [14–28] and custom scripts that connect different parts of the workflow, listed on Table 2.

**Table 1.** Comparison of features between Metaphor and state-of-the-art GRM workflows as listed by [29]. Data adapted to include Metaphor.

| Features | Metaphor v1.7.3 | ATLAS [30] | MetaWRAP [31] | nf-core/mag [32] | MAGNETO [29] |
| --- | --- | --- | --- | --- | --- |
| Preprocessing |  |  |  |  |  |
| Reads trimming | ✓ | ✓ | ✓ | ✓ | ✓ |
| Contamination | ✓ | ✓ | ✓ | ✓ | ✓ |
| Assembly |  |  |  |  |  |
| Coassembly possible | ✓ |  | ✓ | ✓ | ✓ |
| Coassembly by groups | ✓ |  |  |  |  |
| Compute sets to coassemble |  |  |  | ✓ |  |
| Assembly evaluation | ✓ |  |  |  |  |
| Binning |  |  |  |  |  |
| Cobinning possible | ✓ |  | ✓ | ✓ | ✓ |
| Multiple binning software | ✓ | ✓ | ✓ |  |  |
| Bin refinement | ✓ | ✓ | ✓ |  |  |
| Bin reassembly |  | ✓ | ✓ |  |  |
| Postprocessing |  |  |  |  |  |
| MAGs quality check | ✓ | ✓ | ✓ | ✓ | ✓ |
| Dereplication step | ✓ | ✓ | ✓ | ✓ | ✓ |
| Genome annotation | ✓ | ✓ | ✓ | ✓ | ✓ |
| Gene catalogue | ✓ |  |  | ✓ | ✓ |
| HTML Report | ✓ | ✓ |  | ✓ | ✓ |
| Reproducibility |  |  |  |  |  |
| Workflow management | ✓ | ✓ |  | ✓ | ✓ |
| Packages Management | ✓ | ✓ |  | ✓ | ✓ |

**Table 2.**
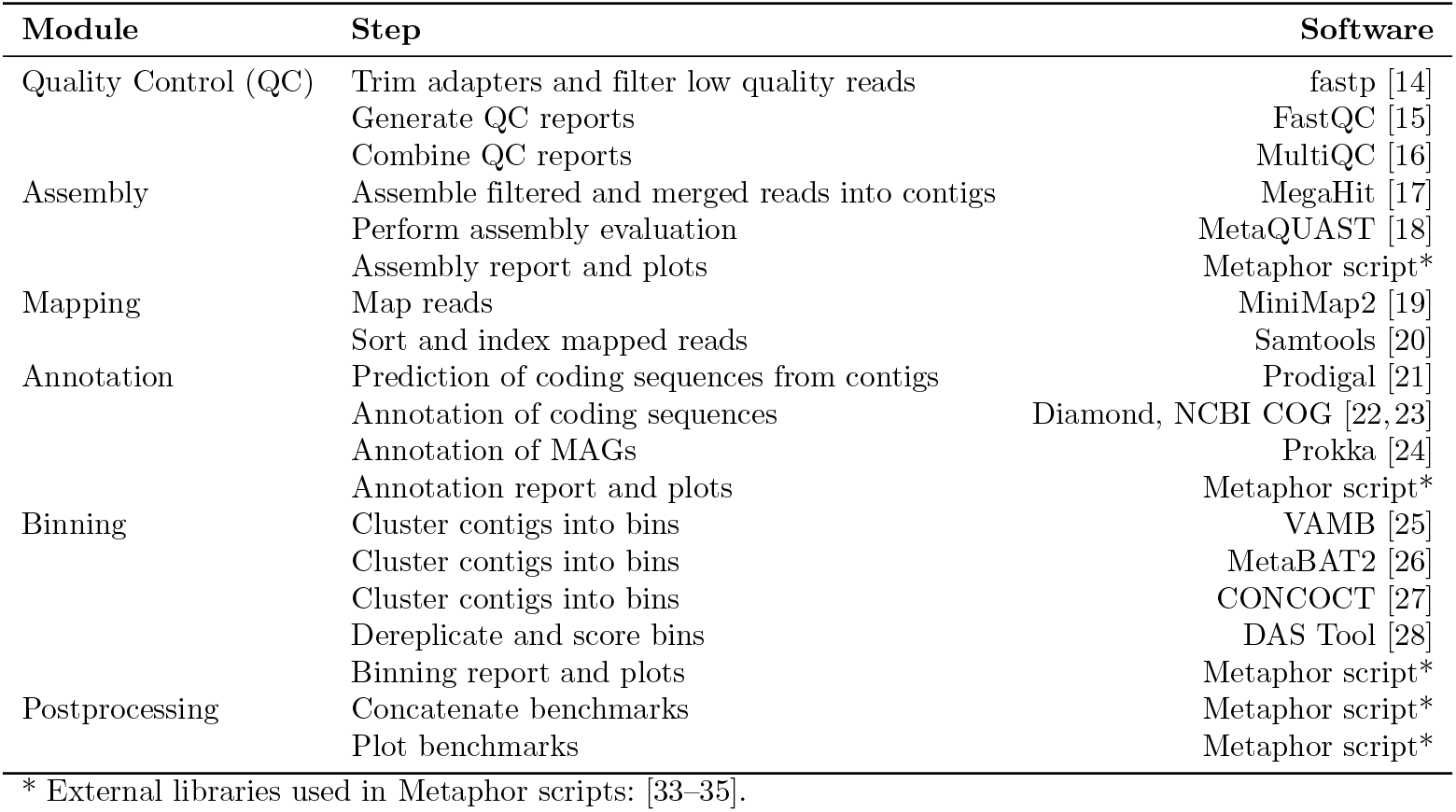
Modules, steps and software used in Metaphor.

The workflow consists of six modules: quality control (QC), assembly, annotation, mapping, binning, and postprocessing. In the QC module, raw sequencing reads are filtered and trimmed. Metagenomic assembly is then performed. Coding sequences are predicted from the assembled contigs and used for functional and taxonomic annotation. The quality-filtered reads are mapped against the contigs, generating coverage statistics employed by the binning algorithms. After binning is complete, bins are refined and dereplicated. Lastly, the postprocessing module renders runtime and memory usage metrics and generates an HTML report. A simplified version of the flow of data between the different modules of the workflow is show on Fig 1.

**Fig 1.**
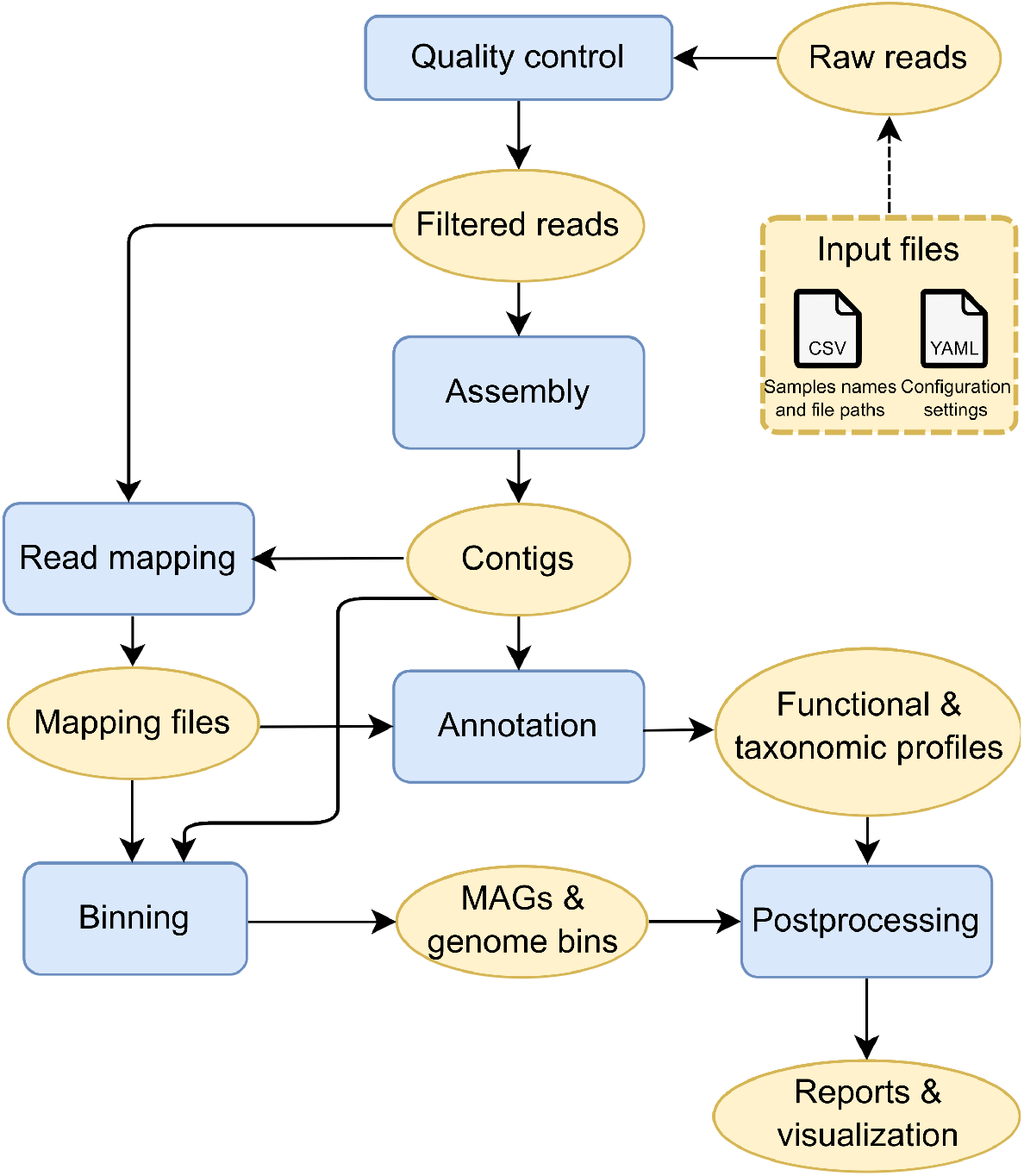
Simplified workflow diagram. Workflow modules are represented by rectangular blue shapes and data files are represented by oval yellow shapes, except for entrypoint files shown in a dashed yellow rectangle. Arrows indicate input and output of data between modules.

The choice of bioinformatics tools was informed by the results of the 2nd Critical Assessment for Metagenome Interpretation (CAMI II) [8, 36], striving for the maximum trade-off between performance, efficiency, and software sustainability. Although the latter is a subjective factor, selecting and streamlining dependencies with regard to code quality, maintenance, and community support is a critical factor when maintaining complex bioinformatics pipelines [6, 37]. Each third-party software (along with its version) is defined in an individual requirements file that is used by Snakemake to create a virtual environment and run that particular step. To facilitate citing these tools, Metaphor packages a bibs/ directory containing all citations in the Bibtext format.

The workflow takes two files as input: a tab-delimited file containing sample names and file paths to the raw reads, and a configuration file in the YAML format, which will set the workflow parameters (see Fig 1). These files can be automatically generated by Metaphor and edited by the user, or created from scratch. The output of Metaphor consists of a directory for each module, further subdivided into the rules within each module. This is described in detail in the documentation [38].

### Assessment on CAMI II synthetic datasets

To demonstrate the functionality of Metaphor, we analysed datasets from CAMI II [8]. All datasets consist of short and long reads generated by simulation of collections of reference genomes [39]). Only short reads were used for each dataset, as Metaphor does not yet support long reads. Specifically, we used the Marine metagenome dataset (identified as ‘marmg’), the Strain Madness dataset (identified as ‘strmg’), and the Human Microbiome dataset, which consists of five sets of samples, each corresponding to a different sampling location in the human body, which were treated as distinct datasets (3). The following strategies were employed for each datasets: single assembly, single binning (‘SASB’), where each sample is individually assembled and binned; single assembly, cobinning (‘SACB’), where each sample is assembled individually and then binned with other samples from the same dataset; coassembly, cobinning (‘CACB’), where all samples from the dataset were assembled and binned together. Table 4 illustrates how this works in practice, in terms of generated output files. Metaphor allows defining multiple groups for coassembly or cobinning to analyse multiple independent datasets with a single execution.

**Table 3.**
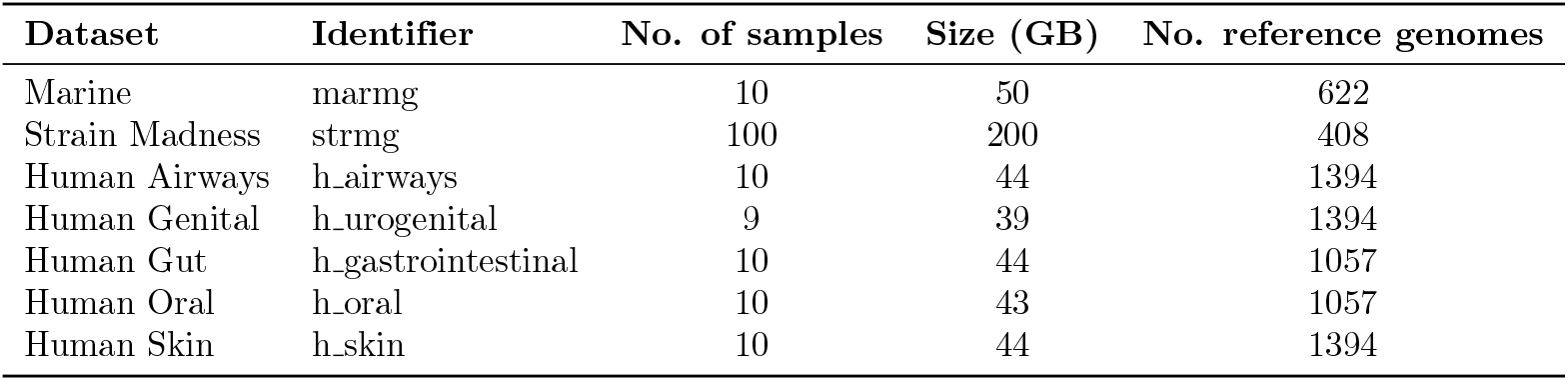
Datasets from CAMI II used to assess the workflow. Columns show the number of samples and size in gigabytes of each dataset, along with the amount of reference genomes used to generate the dataset

**Table 4.**
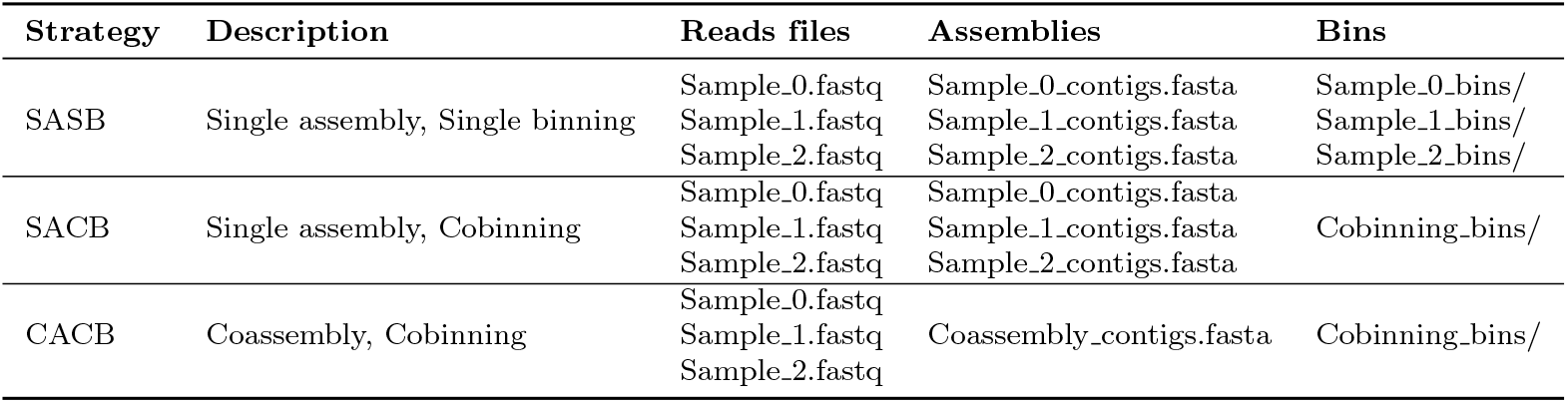
Output files for each strategy. If only one dataset/group is being analysed, assembly and binning results are named as “Coassembly” and “Cobinning” respectively. If multiple datasets/groups are used, the results are named according to the group/dataset’s name.

In order to assess the effect of different assembly strategies, we used MetaQUAST [18] to compare the assemblies generated by the workflow with the collections of reference genomes. For the different binning strategies, we compared metrics obtained from DAS Tool, the software used for dereplicating and evaluating genome bins, after a second round of dereplication with dRep [40]. This is because data generated with the SASB strategy will likely results in redundant bins, as for that strategy there is no dereplication between samples and since samples within a dataset have similar composition, it is likely that a genome bin can be generated repeatedly by different samples. dRep performs dereplication based on the Average Nucleotide Identity between genomes, a metric which has been consistently used as a proxy to differentiate taxonomy at the species and strain levels [41]. dRep was run with default clustering parameters, and without any length, completeness, or contamination cutoffs. We used Spartan [42], the High Performance Computing (HPC) system at The University of Melbourne to run the pipeline. Jobs were dispatched to nodes with the SLURM scheduler, using up to 64 processors and 300 GB RAM per node.

## Results and Discussion

After running Metaphor on the CAMI II Marine, Strain Madness and Human Microbiome datasets, we illustrate the different outputs generated by the workflow, and compare the effects of different assembly and binning strategies on workflow performance.

### Reconstruction of metagenome-assembled genomes

Metaphor produces genome bins generated with three tools: Vamb, MetaBAT2 and CONCOCT [25–27] that are refined with DAS Tool [28]. The input for each binning tool differs slightly, but they all rely on the catalogue of contigs obtained from the assembly and the coverage files obtained from the read mapping module (see Fig 1). A report is generated for each of the binning groups (only one is generated if cobinning is performed), which highlights three key metrics: completeness, redundancy, and bin score. The first two metrics are calculated by the presence/absence of single-copy genes, and the latter is a function of the former two. Plots generated by an example report are shown in Fig 2. It is possible to compare the performance of the different binning software and obtain the proportion of bins above a specified particular quality threshold based on the bin score. The source table for the report is provided, so that users can generate custom reports and inspect specific individual bins. Bins that pass the quality threshold are stored in individual FASTA files, so they can easily be used for downstream analyses with tools such as CheckM or GTDB-Tk [43, 44]. We chose not to include these software in the workflow as they rely on fairly large reference databases and/or contain several different steps that are dependent on third-party software, which would affect Metaphor’s portability. Bin collections generated with Metaphor can be imported into the Anvi’o along with coverage data (BAM files), so users can use the interactive interface of Anvi’o to examine the bins.

**Fig 2.**
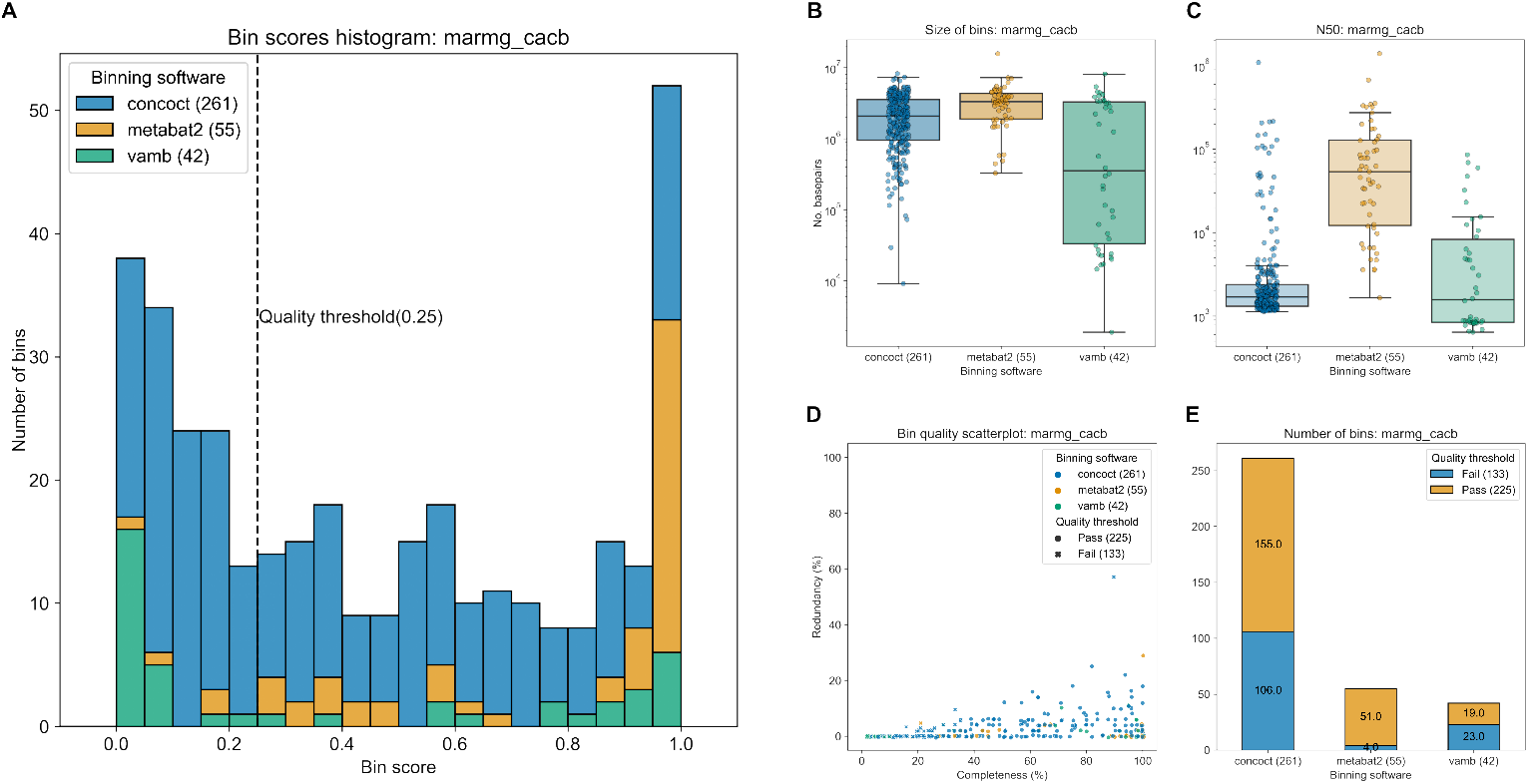
Binning report generated by Metaphor for the CAMI II Marine metagenome dataset processed with the ‘CACB’ (coassembly, cobinning) setting. Panel **A** shows a stacked histogram of the distribution of bin scores, with the defined quality threshold highlighted as a dashed line. Panels **B** and **C** show, respectively, the size (in base pairs) and N50 of bins. The Y-axis is in log-scale. Panel **D** shows a scatterplot of completeness and redundancy for each bin. Colours indicate the tool used to generate the bin, and the symbols indicate whether that bin passed or failed the bin score quality threshold (corresponding to the same value in the dashed line of Panel **A**). Panel **E** shows how many bins passed or failed the quality threshold for each binning tool.

### Contig-level taxonomic and functional profiling

To facilitate quantitative metagenomics applications, Metaphor’s annotation module generates contig-level functional and taxonomic profiles based on the NCBI COG database [23]. These are obtained by predicting coding sequences with Prodigal and then aligning the resulting amino acid files with Diamond [21, 22] in the “iterative” mode. This setting performs repeated rounds of alignment, with an increasing degree of sensitivity when no hits are detected in the previous round. Abundances for each feature are calculated based on the coverage of all coding sequences which align to that feature. Fig 3 illustrates the profile visualisations offered by Metaphor: a heatmap of COG categories for the functional profile and a stacked barplot for the most abundant taxa (for the latter, one plot is generated for each taxonomic rank). The annotation module outputs count tables with both absolute and relative abundance values of taxa and functional categories, and may be directly imported by downstream statistical toolkits such as MixOmics or PhyloSeq [10, 11].

**Fig 3.**
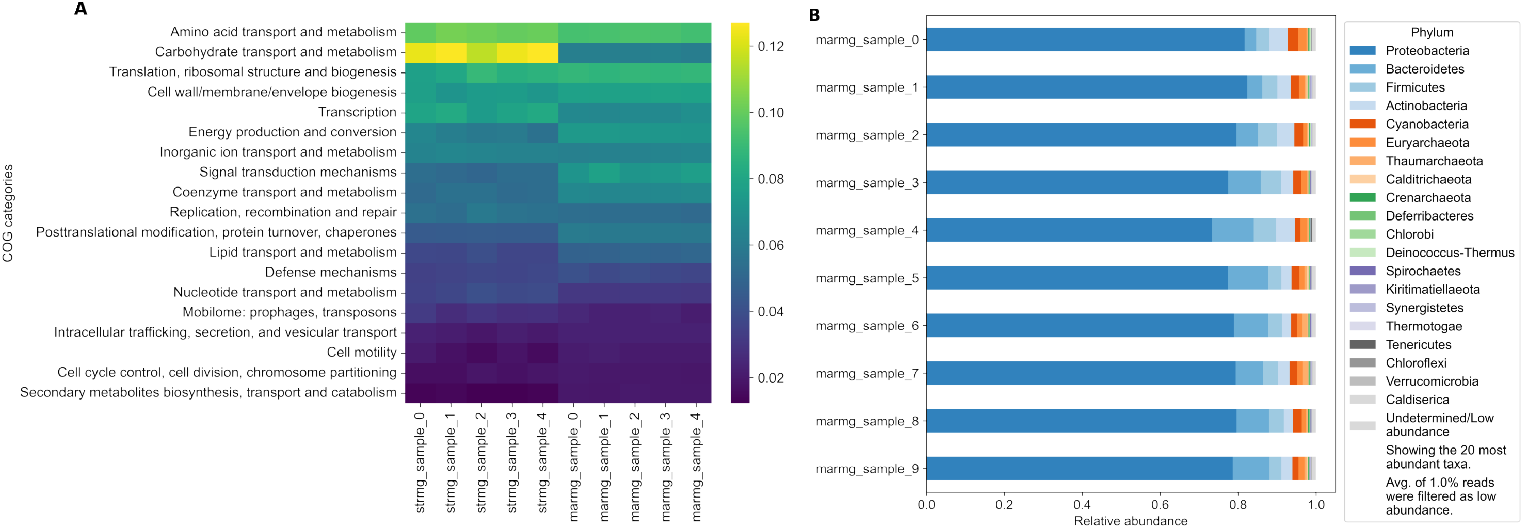
Annotation plots generated by Metaphor on the Strain Madness (‘strmg’) dataset and the Marine dataset (‘marmg’). Panel **A** displays the functional profile as a heatmap of the relative abundance of functional COG categories (Y-axis) across samples (X-axis) for five samples from Strain Madness and Marine datasets. Panel **B** displays the taxonomic profile of the Marine dataset as a stacked barplot of relative abundance of taxa. In this case, the phylum rank was used, but Metaphor generates this for the most common taxonomic ranks (phylum, class, order, family, genus, species). The number of abundant taxa can be easily adjusted in the workflow settings. For both taxonomic and functional profiles, abundance of each feature is calculated from coverage values for each gene.

### Quality control and performance metrics

Additional outputs produced by Metaphor include the quality control reports from the fastp and FastQC tools, with a summary of FastQC outputs being produced by MultiQC [14–16]. A simple report is produced by the assembly module with sequence statistics of the assembled contigs (*e.g*. N50, number of contigs, total and mean length of contigs), and performance metrics. At the end of the workflow execution, the postprocessing module generates figures obtained from the “benchmark” files provided by Snakemake. These files contain process information such as runtime and memory consumption. Metaphor plots these metrics in two ways: total per rule and per-sample mean (Fig 4) as some rules run only once across all samples, while other rules run per sample. These plots help identify computational bottlenecks and assess whether computing resources are adequate.

**Fig 4.**
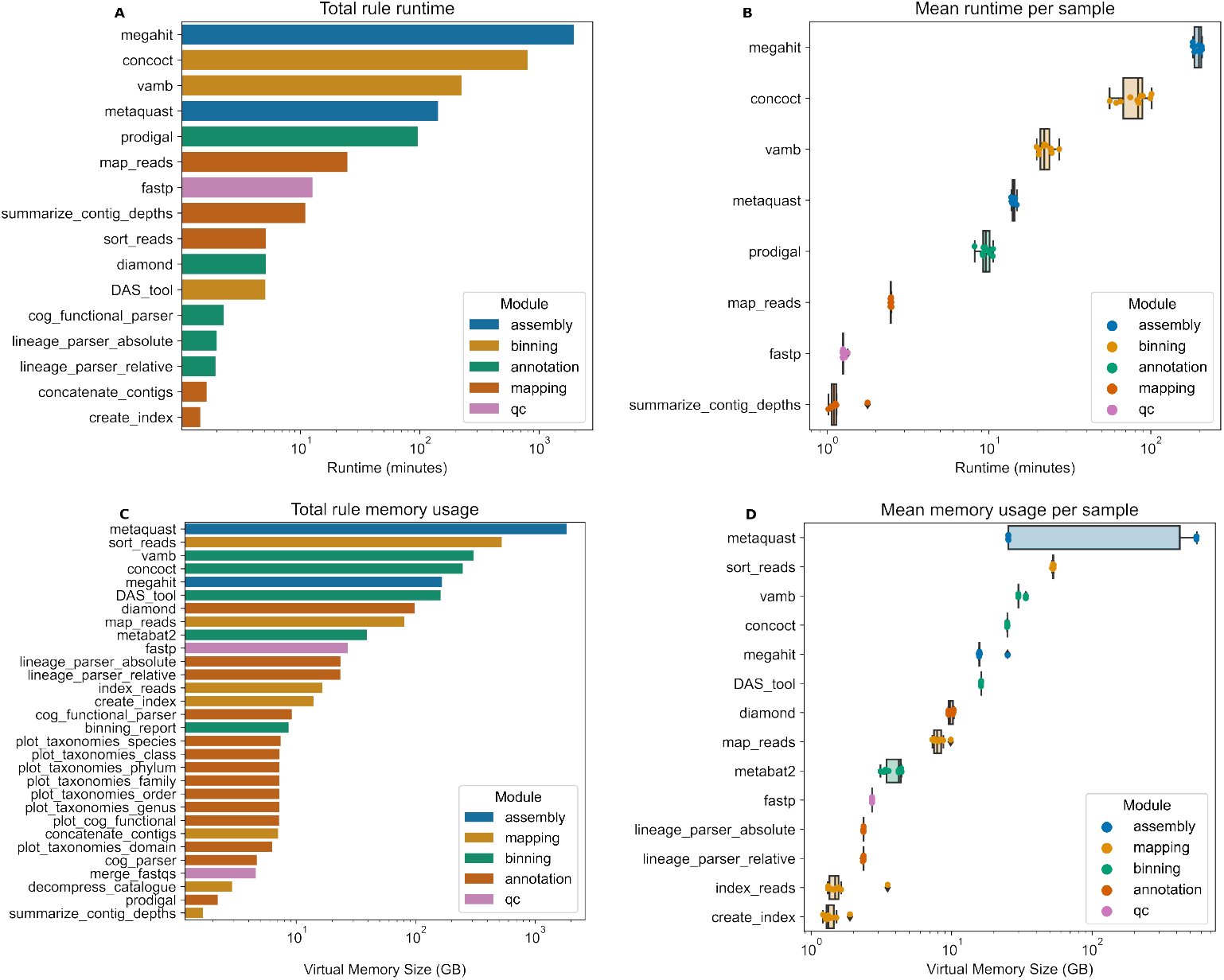
Performance metrics report generated by Metaphor on the Marine dataset processed with the SASB strategy. Total runtime per rule (**A**), mean runtime per sample (**B**), total memory usage per rule (**C**), and mean memory usage per sample (**D**). X-axis is in log format. Cutoffs are applied to omit rules with short runtime or low memory usage. Colours indicate the workflow module of each rule.

### Assembly and binning strategies

The effects of distinct assembly and binning strategies on the final output of metagenomic workflows are highly dependent on the data source and research context [8]. As such, the choice of individual or group assembly and binning can only be assessed a posteriori. We compared three different strategies: single assembly and single binning (‘SASB’), single assembly and cobinning (‘SACB’), and coassembly and cobinning (‘CACB’), see Table 4 and Section ‘Assessment on CAMI II synthetic datasets’ for details. For assembly, we used the five different groups in the Human Microbiome dataset along with the Strain Madness and Marine Datasets. We only used the latter two datasets for the binning assessment.

We used six metrics to evaluate assembly performance: percentage of recovered genome fraction, size of the largest contig, duplication ratio, length of misassembled contigs, number of misassemblies, and number of mismatches per 100 thousand base pairs. High values for the first two metrics and low values for the last four indicate better performance. We observed a general trade-off between assembly completeness (represented by the first two metrics), and the number of errors in the assembly (represented by the last four metrics) (Figure S1). In most datasets, assemblies were more complete and contiguous, albeit with more errors when the Coassembly strategy was used. The exception was the Strain Madness (‘strmg’) dataset, for which the Individual assembly was more complete and contiguous, albeit with more errors. This may be attributed to the high degree of strain/intraspecies diversity in that dataset [8]. A high degree of similarity between the related genomes likely confounds assembly algorithms, and pooling samples together may aggravate this effect [5].

To evaluate differences between binning strategies, we compared the number and quality of bins after refinement with DAS Tool. Bins generated with each approach were further dereplicated with dRep [40]. This is because the SASB strategy generates a set of bins for each sample, and datasets with similar composition will likely generate redundant bins, as there is no dereplication of bins between samples. Results varied significantly between the Marine and Strain Madness datasets. In both datasets, the mean bin score was the highest for the CACB strategy (S3 Fig). However, in the Strain Madness dataset, CACB produced a significantly lower number of bins (33 compared with 259 and 215 generated with SASB and SACB), which did not occur in the Marine dataset.

Since the binning performance is assessed as a proxy of the combination of quantity and quality of generated bins, rather than only one metric or the other, we calculated the cumulative bin score (the sum of scores of all bins) and the number of bins above an increasing score threshold, shown on Fig 6. The higher the threshold, the more significant the differences between the cumulative scores, as only bins with the highest quality compose the score. For the Marine dataset, we observed a higher score and a larger number of bins in the CACB strategy and the exact opposite in the Strain Madness dataset. In both datasets, there was a clear difference between SASB before dereplication and the other strategies, confirming that several highly similar samples produce redundant bins. That difference was also present in the SACB strategy, albeit not so pronounced (see S4 Fig for the comparison of dereplicated and non-dereplicated data). This suggests that for both of these strategies, further dereplication is recommended [5]. Although the Strain Madness dataset shows fewer bins generated with CACB, the cumulative bin score for that strategy remained similar to SACB and SASB above the 0.8 score threshold, since there are fewer bins with a score lower than that. In that same dataset, SASB showed the best performance, although differences were small above the 0.8 threshold. In the Marine dataset, there were more pronounced differences between strategies. CACB produced the larger quantity and higher cumulative score of bins, followed by SASB and SACB.

**Fig 5.**
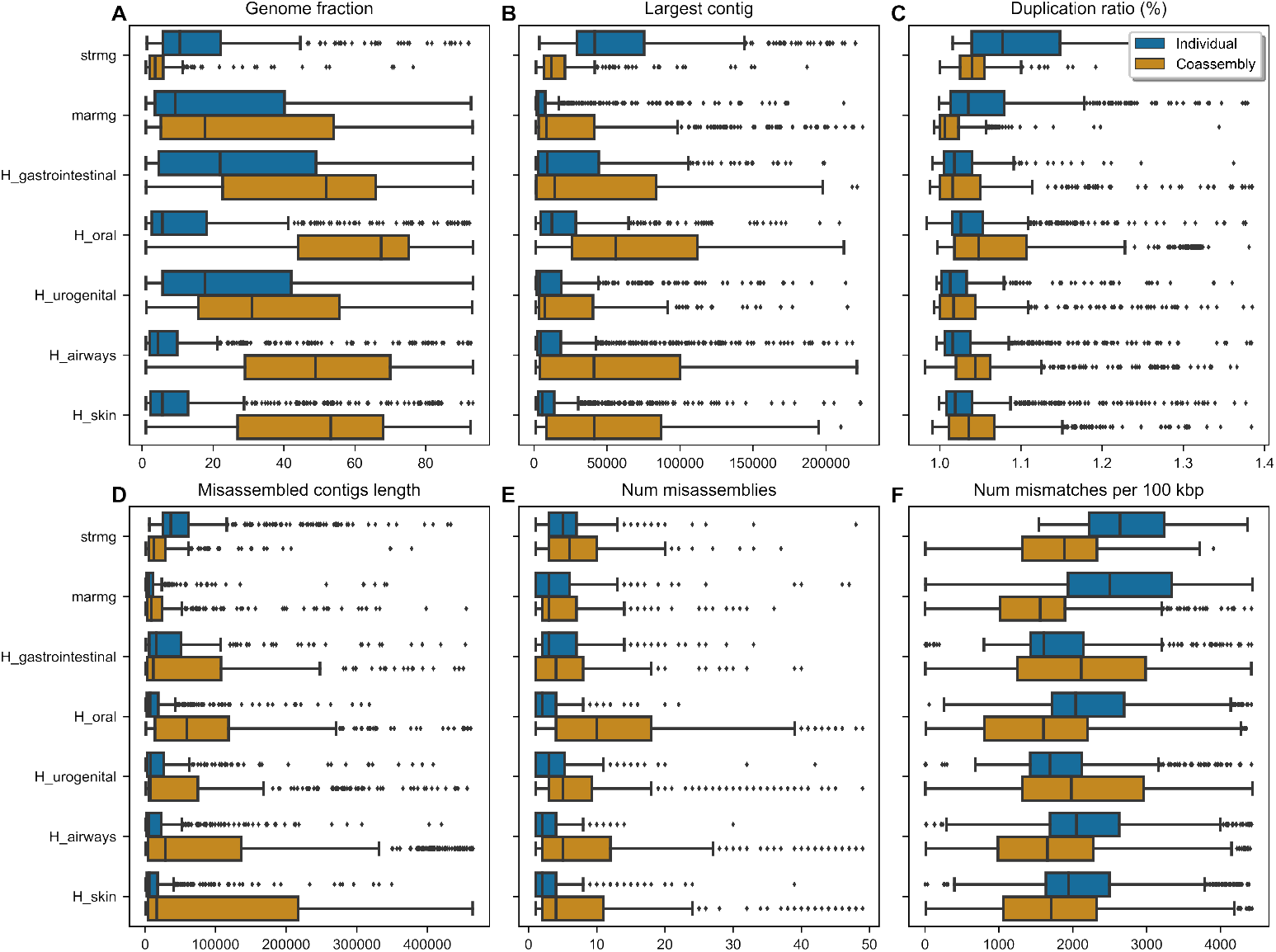
Differences between assembly strategies for each dataset. Each data point corresponds to a reference genome evaluated with the MetaQUAST tool. Data points above the 98th percentile were classified as outliers and removed from the figure to improve visualisation. See S1 Fig for the full data. The title at the top of each panel indicates the plotted metric. Panels **A** and **C** show percentages along the X-axis, while the remainder show absolute values.

**Fig 6.**
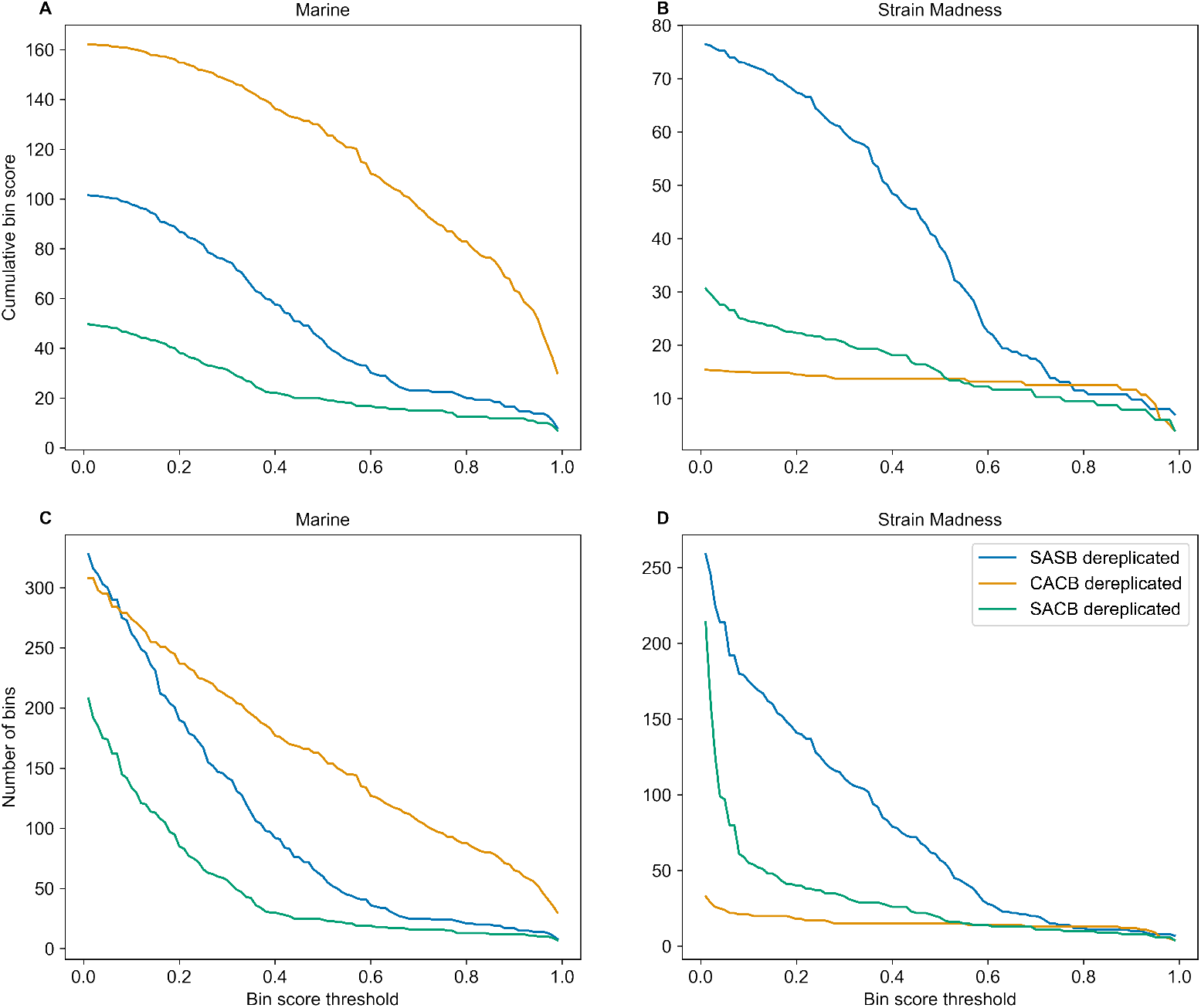
Cumulative bin score and number of bins between binning strategies for the Marine and Strain Madness datasets. Lines show the cumulative bin score (**A** and **B**) and number of bins (**C** and **D**) along the Y-axis, for bins above a certain score threshold (X-axis). Left column shows Marine dataset, and right column shows Strain Madness dataset.

In summary, our assessment of different assembly and binning strategies indicate that, for most metagenomic analysis scenarios, coassembly followed by cobinning is samples are sourced from a similar environment or population, except when there is a high level of intraspecies/strain-level diversity across samples, like in the Strain Madness dataset. In that scenario, single assembly followed by single binning is preferred, followed by dereplication of bins between samples. There is, however, a trade-off, as computational requirements are higher for the pooled strategies. Coassembly resulted in higher genome recovery fractions and larger contigs, although usually at the expense of a higher number of misassemblies and higher duplication ratio. When combining coassembly with cobinning, there is a remarkable improvement in the quantity and quality of bins generated for a diverse dataset (represented by the Marine dataset), where the difference was negligible in the Strain Madness dataset.

## Availability and Future Directions

Metaphor is available through Bioconda [45], a popular repository of bioinformatics software. It can be installed with a single command from the conda package manager [46] or from source using pip, the Python package manager. The installation of all third-party software used by Metaphor is handled automatically by Snakemake and conda. Metaphor is developed with documented best practices in workflow development [6, 47], striving for reproducibility and transparency of its results. Documentation for Metaphor is available at https://metaphor-workflow.readthedocs.io/.

The workflow may be extended to support downstream tools such as GTDB-Tk and CheckM, and a new functionality for the identification of eukaryotic and viral contigs and bins. The annotation module can also be improved to facilitate the use of custom reference databases. In addition, Metaphor would benefit from new third-party software to facilitate the generation of non-prokaryotic bins in the near future.

## Acknowledgments

Metaphor benefited strongly from experience gained developing MetaGenePipe (in press [48]), a Cromwell-based workflow for assembly and annotation of metagenomic contigs. VWS is funded by a Melbourne Research Scholarship from The University of Melbourne. This research was supported by The University of Melbourne’s Research Computing Services and the Petascale Campus Initiative. We thank Francesco Ricci and Uthpala Pushpakumara for providing datasets for early trials of Metaphor, and colleagues from the Lê Cao lab for sharing their feedback.

## Supplementary Material

**S1 Fig.**
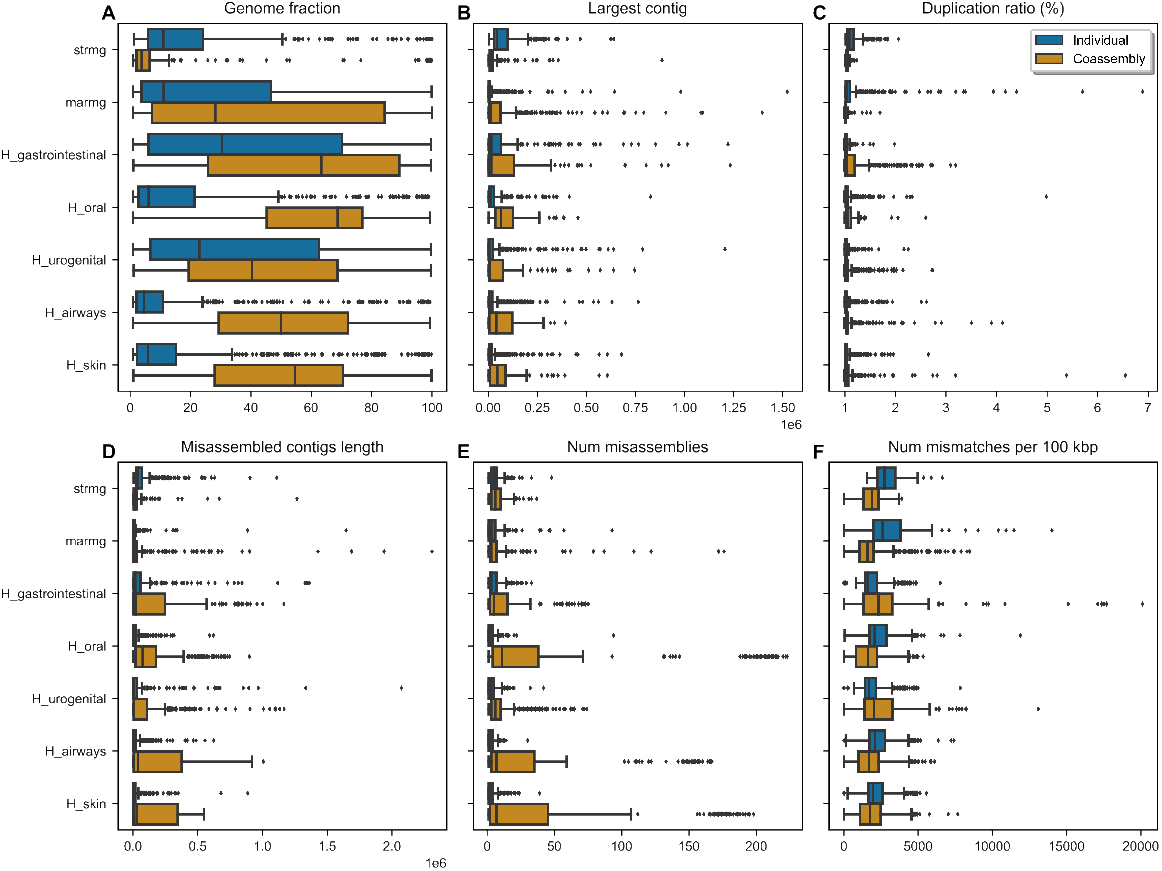
Differences between assembly strategies across datasets. Same data as Fig 5, but including outliers.

**S2 Fig.**
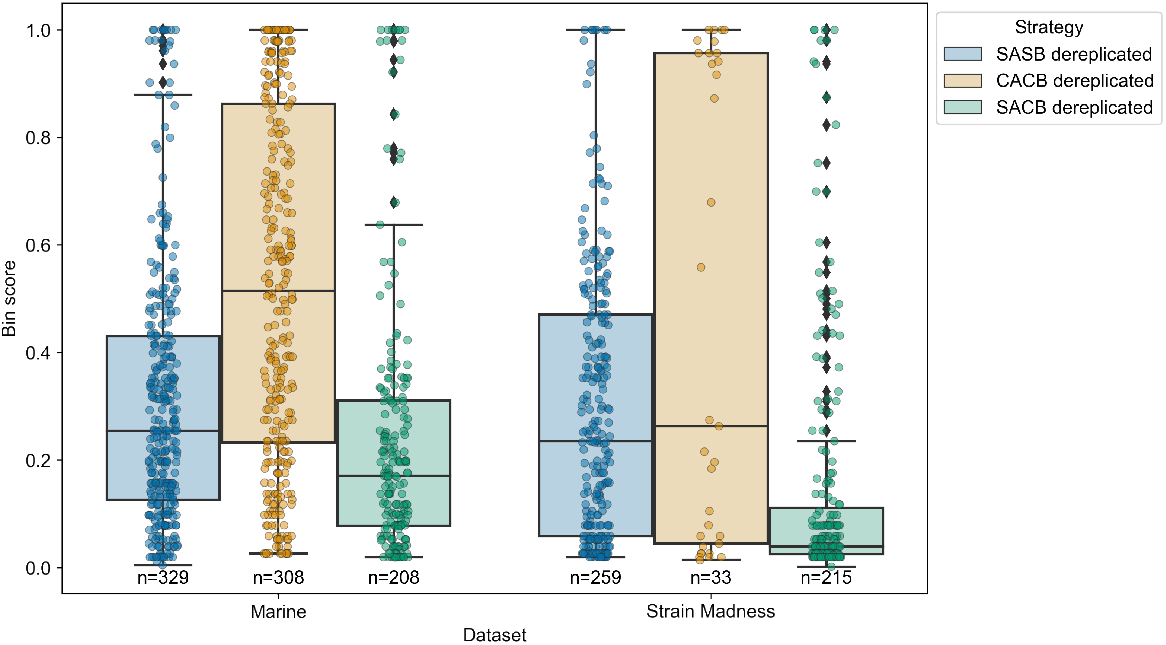
Boxplot of bin scores across different strategies. Each data point is a genome bin, and Y-axis depicts bin scores from 0 to 1. Columns separate datasets, and colours represent different strategies. Numbers underneath each bar show the number of data points for that bar. Bins sets were dereplicated with dRep.

**S3 Fig.**
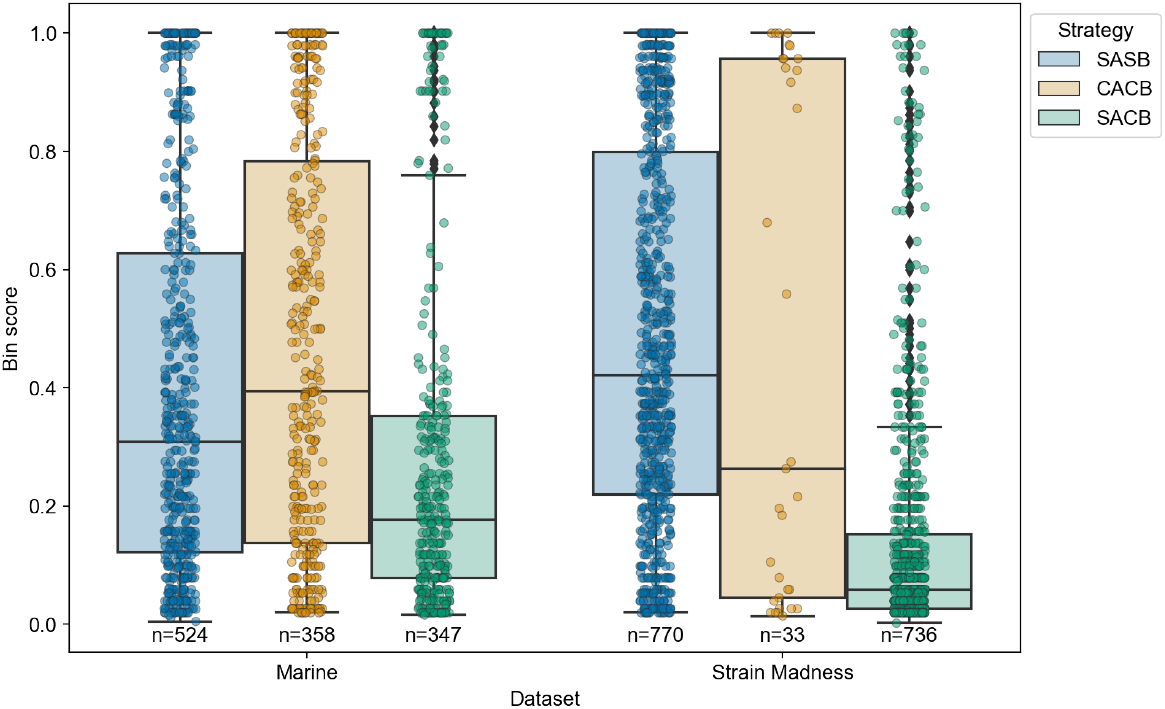
Boxplot of bin scores across different strategies for non-dereplicated data. Same as S2 Fig, but with non-dereplicated data. Each data point is a genome bin, and Y-axis depicts bin scores from 0 to 1. Columns separate datasets, and colours represent different strategies. Numbers underneath each bar show the number of data points for that bar.

**S4 Fig.**
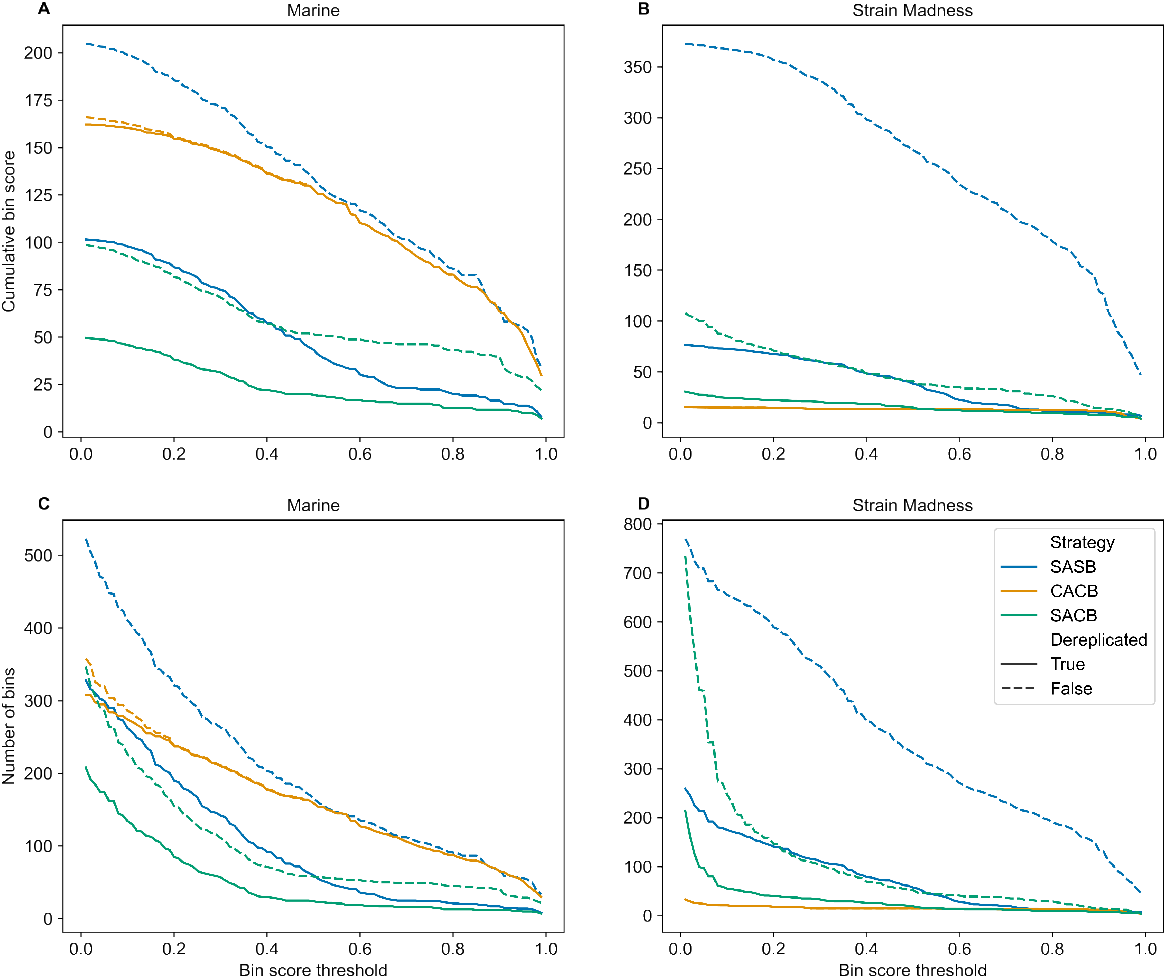
Cumulative bin score and number of bins between binning strategies for the Marine and Strain Madness datasets. Solid lines show the same data as Fig 6, and dashed lines show data based on bins prior to dereplication with dRep.

## Notes

https://github.com/vinisalazar/metaphor

### Competing Interest Statement

The authors have declared no competing interest.

